# Integrating knowledge, omics and AI to develop patient-specific virtual avatars

**DOI:** 10.1101/2024.11.07.622508

**Authors:** Amel Bekkar, Luca Santuari, Ioannis Xenarios, Bulak Arpat

## Abstract

We propose a method for creating personalized regulatory networks, enabling the development of virtual avatars for cancer patients, with patient-derived xenograft (PDX) models as a test case. Starting from a Prior Knowledge Network (PKN) based on the hallmarks of cancer, we constructed gene networks that are contextualized to each sample by integrating sample-specific gene expression data. These networks were optimized using a genetic algorithm to align with individual molecular profiles, focusing on key cancer-related processes. Following network optimization, we employed Graph Convolutional Networks (GCNs) to classify samples based the structures and interactions of their individualized network models and molecular profiles. This personalized approach provides insights into drug responses and helps predict treatment outcomes, offering a path toward more targeted cancer therapies.

**Author summary:** Cancer treatment can be more effective when therapies are personalized to each patients unique molecular profile. In this study, we introduce a method to create virtual avatars of cancer patients by personalizing regulatory networks using patient-derived xenograft (PDX) models as a proof of concept. Starting from known cancer hallmarks, we developed individualized gene networks for each sample by leveraging their specific gene expression data. These networks were refined with an optimization process to match the distinct molecular characteristics of each sample. By applying advanced machine learning, specifically Graph Convolutional Networks (GCNs), we classified these personalized models to better understand likely drug responses and predict treatment outcomes. This approach brings us closer to tailoring cancer therapies to individual patients, potentially improving treatment success by targeting key cancer pathways unique to each person.

## Introduction

Integrating knowledge and omics data to model gene regulatory networks has long been a pivotal approach in understanding the complexities of biological systems. This multifaceted strategy not only aids in deciphering mechanisms of action at a molecular level but also enhances our ability to make precise predictions about cellular behavior. By leveraging high-throughput data from omics technologies alongside established biological knowledge, researchers can construct comprehensive models that reveal intricate regulatory relationships and interactions. [1–4].

The development of these models is crucial for several reasons. Firstly, they can lead to the identification of novel biomarkers and therapeutic targets. Secondly, they enable the prediction of cellular responses to various stimuli, including drugs and environmental changes. Lastly, these models contribute to the understanding of disease mechanisms, facilitating the discovery of new treatment strategies and the improvement of existing ones [5].

An emerging and particularly promising application of this approach is the creation of personalized models, referred to as *in silico* avatars. These avatars simulate individual patient biology, allowing for the prediction of disease progression and response to treatment on a highly personalized level taking into account the subtle biological differences between people [6]. This capability is instrumental in picking the right target for treatment and determining which patients are most likely to benefit. Additionally, avatars can aid in the development of drug combination strategies, biomarker discovery, and the prediction of resistance, providing critical insights for novel treatment approaches [7]. Simulation will help experimental work and clinical trials by carrying only molecules and targets with the best chances of success and finding the subset of patients who would get the most benefit from a treatment.

Despite the significant progress in this field, challenges remain. Addressing the inherent complexity and high-dimensionality of gene regulatory networks (GRNs) requires sophisticated computational methods and algorithms capable of handling large datasets, integrating diverse omics data and extracting meaningful patterns [8]. Recent advancements in computational methods, particularly in machine learning and graph neural networks (GNNs), have significantly enhanced our ability to model and analyze these networks. GNNs, by leveraging graph structures to capture the intricate relationships between genes, proteins, and other biological entities, have improved the scalability and accuracy of predictions related to gene regulation and potential interventions [9].

This paper aims first to address these challenges in two ways: first, by presenting novel methodologies for the integration and analysis, highlighting the potential of integrative approaches and advanced graph-based computational methods; and second, by showcasing real-world applications of these technologies, using *in silico* avatars to model baseline tumors and response to treatment scenarios as a case study.

### 0.1 Current state of the art

We believe that logic-based models are particularly well-suited for virtual avatars due to their ability to model large biological networks without requiring detailed kinetic data. This approach allows researchers to simulate complex cellular processes and predict the outcomes of various perturbations, such as drug treatments. For example, the study by Singh et al. [4] presents a comprehensive Boolean model of rheumatoid arthritis (RA). This model simulates RA fibroblast-like synoviocytes (RA-FLS) behavior which helps to predict drug synergies, identify novel therapeutic targets, and simulate the effects of drug combinations, offering a powerful tool for RA treatment optimization and drug repurposing. Moreover, the study by Tsirvouli et al. [10] uses a Boolean network model combined with stochastic simulations to explore the molecular mechanisms underlying psoriasis, allowing for the identification of potential drug targets by simulating various interventions within a complex network of cellular interactions. In addition, the work by Singh et al [11] employs a logic-based framework to predict effective therapeutic strategies for melanoma by targeting E2F1-driven pathways, showcasing the potential for identifying novel drug combinations and repurposing existing drugs to impede melanoma progression. Together, these studies exemplify the versatility of logic-based models in advancing therapeutic discoveries across diverse disease contexts.

Logic-based models have been notably used in cancer research for constructing patient-specific models. For instance, Montagut et al. [2] developed Boolean models for prostate cancer based on patient-specific molecular data from The Cancer Genome Atlas (TCGA) resource. These models simulate the impact of various drug interventions, providing personalized treatment strategies based on the patient’s unique molecular profile. The models were also validated experimentally on prostate cancer cell lines, demonstrating their ability to guide precise interventions for different patients. Similarly, in a study by Eduati et al. [1], the authors present a methodology for creating personalized logic models of cancer signaling pathways, tailored using high-throughput data from cancer biopsies. The main goal is to prioritize and predict effective combination therapies for individual patients.

We think that GNNs are a valuable complement to Boolean frameworks, particularly for integrating multi-modal data and managing large collections of patient-specific networks. GNNs have rapidly advanced in recent years, becoming highly effective in tasks like gene regulatory network (GRN) inference, which has been a long-standing challenge in the field [12]. Wu et al. [13] have shown how GNNs can capture complex gene interactions through approaches like multi-view hierarchical hypergraph.

Additionally, models like MOGONET, designed for multi-omics classification and biomarkers identification [14] uses GCNs to integrate gene expression, DNA methylation, and miRNA expression data, providing a robust platform for analyzing complex biological data and constructing GRNs. This method and others have been reviewed by Wekesa and Kimweley [15].

Some models combine GNNs with multi-omics data (e.g., genomics, transcriptomics) alongside prior knowledge networks like protein-protein interactions. The MPK-GNN framework is an example of a system that applies prior knowledge to guide the integration of omics data into a GNN model.

The study by Roohani et al. [16] introduced GEARS, a geometric deep learning method designed to predict the transcriptional responses to both single and multigene perturbations. GEARS leverages single-cell RNA sequencing data and a knowledge graph of gene-gene relationships to model how genes interact under various perturbations. A key feature of GEARS is its ability to predict the outcomes of novel, untested gene combinations, making it a valuable tool for guiding experimental design.

Similarly, Li et al. [17] introduced PINNACLE, a geometric deep learning approach specifically designed to generate context-aware protein representations. The model integrates data from a multi-organ single-cell atlas, leveraging protein interaction network. One of PINNACLEs key innovations is its ability to adjust its outputs based on biological context, such as the specific tissue or cell type in which a protein is active.

Finally, Dorier et al. [18] developed Optimusqual, a method for reconstructing regulatory networks using genetic algorithms and prior biological knowledge. This enables the creation of patient-specific networks by tailoring the model to individual biological profiles. Integrating these patient-derived networks with GNNs could significantly enhance their predictive power. GNNs, with their ability to model complex relationships and topologies, would amplify the accuracy of patient classification and improve predictions related to treatment response. By combining the structured biological insights from Boolean regulatory networks with the scalability and learning capabilities of GNNs, this approach could lead to more precise and personalized medical predictions, facilitating the discovery of novel therapeutic targets and biomarkers (fig 1).

**Fig 1.**
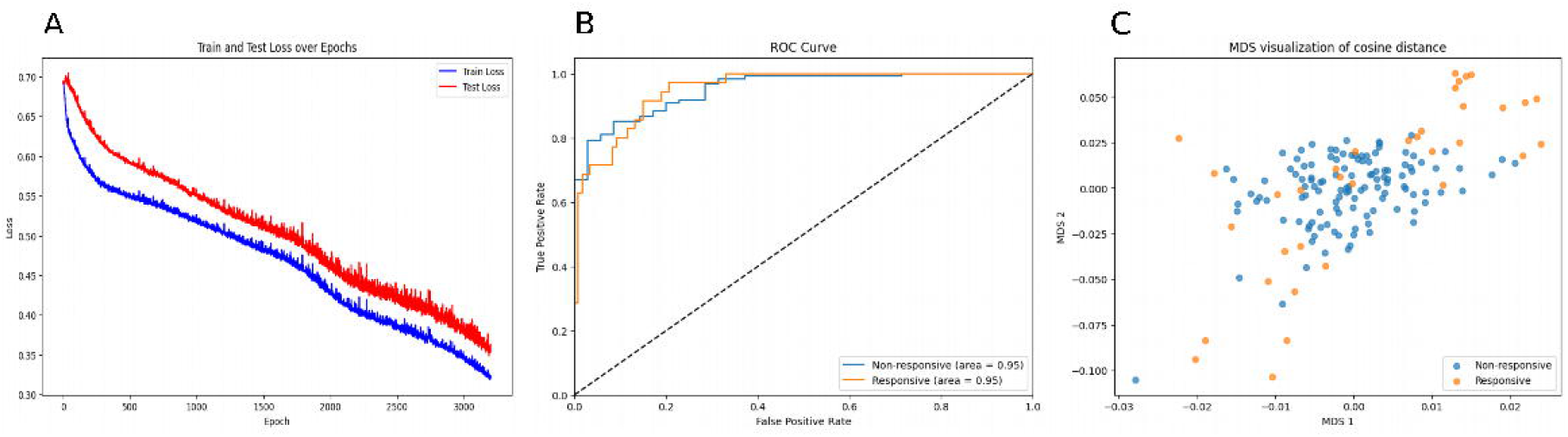
Schematic representation of Virtual Avatar-based response prediction. This figure illustrates the process of using virtual avatars to predict patient-specific treatment responses. The workflow begins with two main sources of data: Prior Knowledge Networks (left) and Patient-Specific Omics Data (right). These data sources are integrated into Virtual Avatars (center), which simulate the patient’s disease progression and potential treatment response. From these avatars, two key outputs are generated: Mechanistic insight: Understanding the underlying biological mechanisms driving the treatment response or resistance; and response prediction: Predicting the likelihood of a patient responding to a specific therapy.

## 1 Results

### 1.1 Building executable patient derived models

As a use case for digital avatars, we evaluated the response of four patient-derived xenograft (PDX) models – two esophageals (ES0001 and ES0002), one lung (LU0001) and one pancreatic (PA0001)– from Crown Bioscience International collection, to an NRG1-Ab [19]. Among these models, only one (ES0001) exhibited a positive response. To understand the differential response, we constructed personalized models for each individual sample by integrating prior knowledge with sample-specific expression data.

#### Prior knowledge network

We initiated the modeling process by assembling a prior knowledge network (PKN), which encompasses known gene regulatory relationships relevant to the hallmarks of cancer [20]. Curated databases and tools, such as MsigDB [21], Signor [22] and Omnipath [23], proved highly useful in constructing the PKN. These resources enabled us to integrate diverse sources of regulatory information, providing a comprehensive foundation for our model that captures the complexity of cancer-related signaling pathways. The resulting network is composed of 1202 nodes and 9045 edges, reflecting the intricate interactions involved in cancer biology.

#### Models personalization using genetic algorithm

For each PDX sample, using its gene expression data we contextualized the prior knowledge network using Optimusqual. This optimization process tailored the network to reflect the unique expression data of each sample. The genetic algorithm iteratively adjusted the network to maximize its alignment with the baseline and the treated expression profiles of the respective sample. To explore the diversity of possible networks that could fit the expression data, we performed 30 replicates of optimization for each sample. However, due to the large size of the PKN, not all optimizations were successful, resulting in a range of 19 to 30 successful replicates per sample. All optimized networks achieved a score higher than 0.9, indicating that the PKN contains sufficient information to explain the gene expression data of each sample (Supplementary fig1 **??**).

**Supplementary Figure 1:**
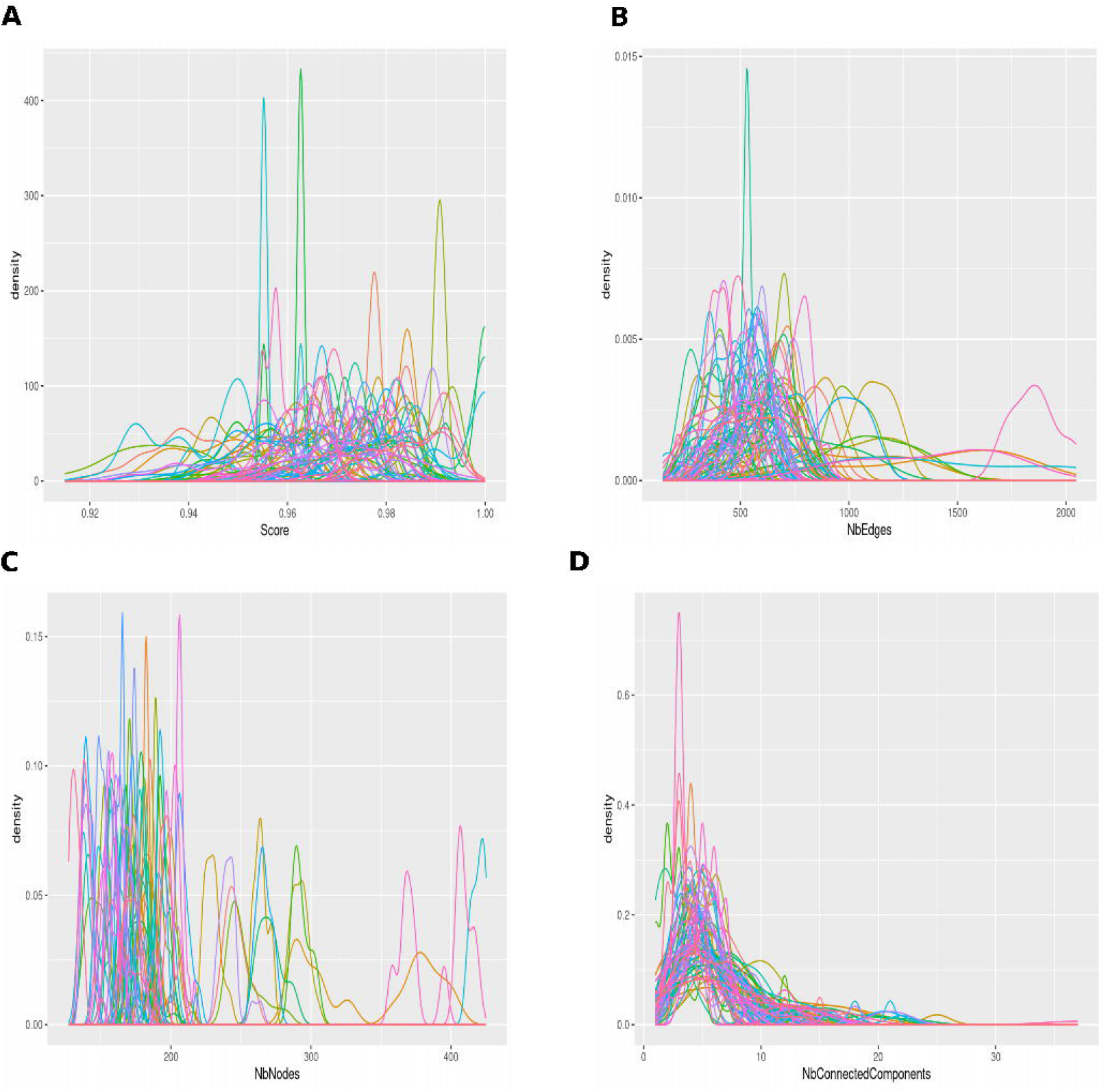
Density distributions of network properties across simulated regulatory networks per sample. (A) Distribution of network scores. (B) Distribution of the number of edges. (C) Distribution of the number of nodes. (D) Distribution of the number of connected components.

#### Boolean modeling and attractor calculation

To further analyze the behavior of these optimized networks, we employed Boolean modeling to calculate the attractors. Attractors in Boolean models represent stable states or patterns of gene expression that the network can adopt.

Attractor analysis revealed distinct patterns between the responsive and non-responsive PDX models (fig 2): The optimized network for the responsive PDX model (ES0001) demonstrated a clear separation between NRG1-Ab treated and non-treated samples. This separation was evident in the attractor profiles, indicating distinct stable states under treatment versus baseline conditions. In contrast, the networks for the non-responsive PDX models did not show such a separation. The attractors profile for NRG1-Ab treated and non-treated samples overlapped significantly, suggesting that the treatment did not induce a notable change in the stable expression states of these models.

**Fig 2.**
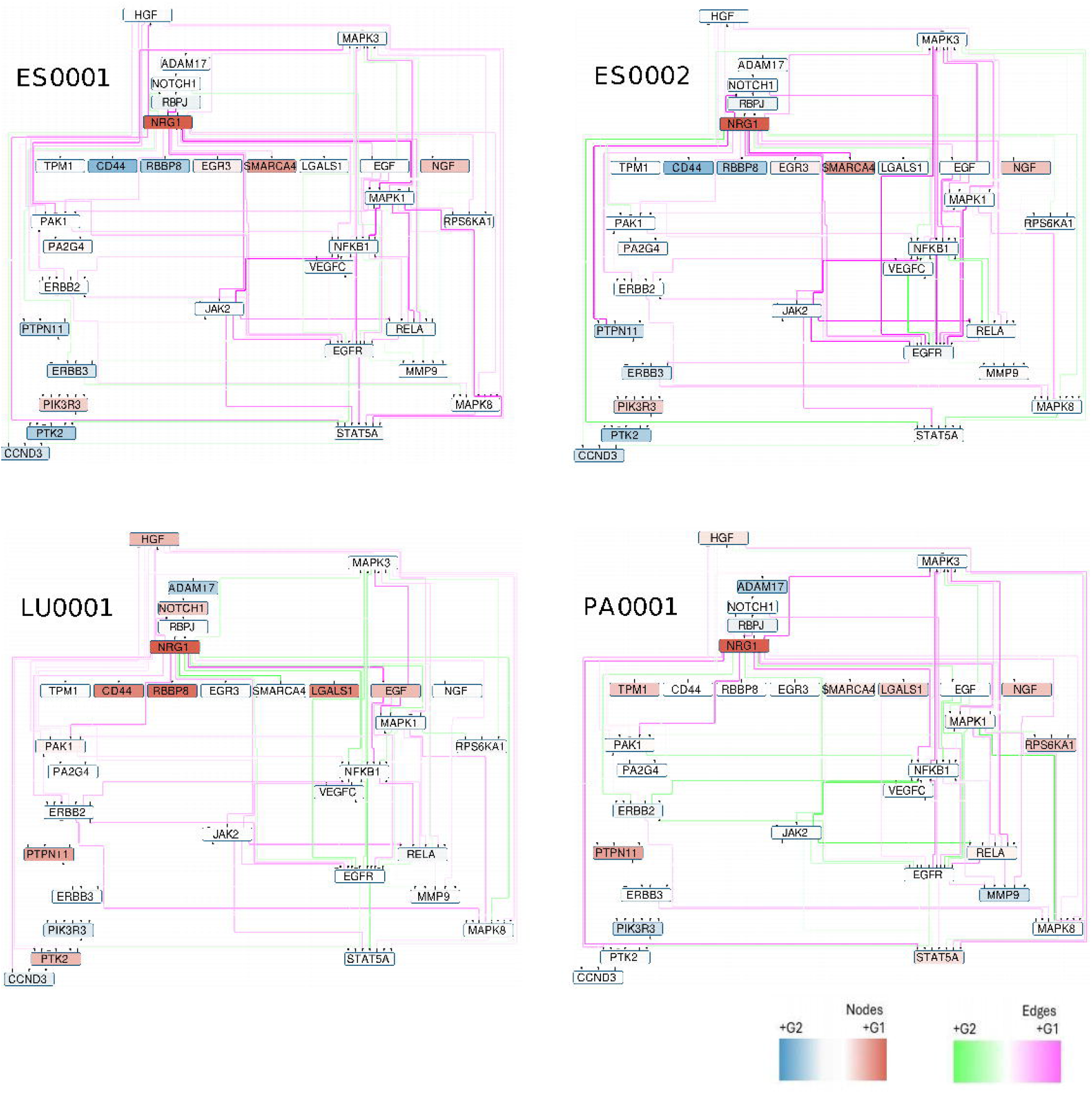
Heatmap representation of attractor states for NRG1-Ab treatment groups. This heatmaps illustrate the attractor states of individual models for four different samples: ES0001 (top-left), ES0002 (top-right), LU0001 (bottom-left) and PA0001(bottom-right) under two treatment conditions: NRG1-Ab- (green) and NRG1-Ab+ (red). Columns: Each column represents a specific node in the gene regulatory network. Rows: Each row represents the median attractor state of an individual sample, which is calculated by first determining the mean attractor state for each replicate model of that sample, then, the median of these mean attractor states is computed across all replicates. The color scale (ranging from white to dark blue) indicates the attractor state values, where:White (0) represents an inactive state.Dark blue (1) represents an active state. On the left side, the color bars denote the treatment group for each sample: Green represents NRG1-Ab- (control) samples.Red represents NRG1-Ab+ samples.

### 1.2 EPDMs to study mechanism of action

To further investigate the mechanisms underlying differential responses to antibody treatment, we analyzed personalized gene regulatory networks, focusing on NRG1 and its surrounding network structure. Specifically, we examined within-group edge frequency in treated and non-treated samples and compared edge frequencies between the responsive and non-responsive groups. We identified significant variations in network motifs when focusing on the regions surrounding NRG1. The differences in edge weights between these groups were particularly notable, suggesting altered structural integrity of non-responsive models compared to the responsive one.

(Fig 3) highlights the distinct differences in network motifs and feedback loops between responsive and non-responsive models. Interestingly, while NRG1 itself was predicted to be inactive in the attractor states under antibody treatment in both groups, the key variations in network structure were located in other regions of the network. This suggests that the differential treatment response may be driven by interactions within the broader regulatory network, rather than solely by NRG1.

**Fig 3.**
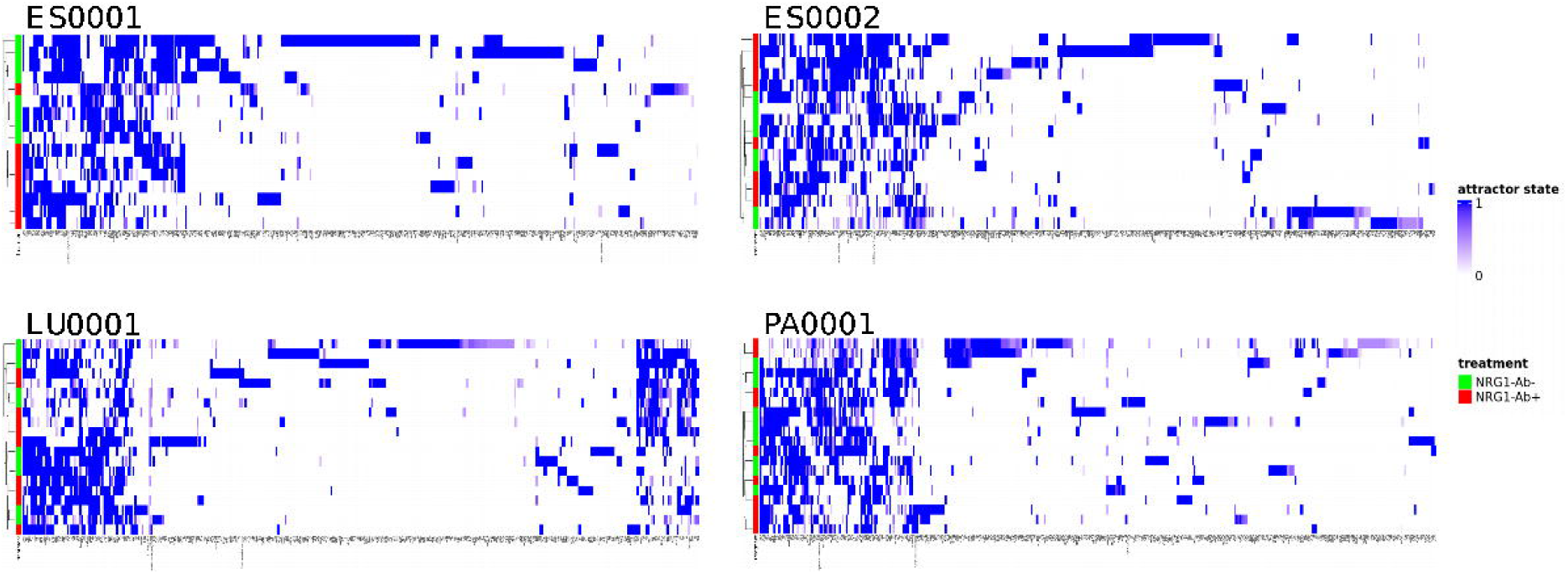
Network representation of NRG1 neighbors in NRG1-Ab Treatment Groups. This figure shows the personalized gene regulatory networks for four different samples (ES0001, ES0002, LU0001, PA0001) under two treatment conditions: NRG1-Ab- (G1) and NRG1-Ab+ (G2). Each network illustrates the nodes and edges representing the gene interactions and their changes between the two groups. Nodes: The color gradient of the nodes represents their differential activity between the two conditions. Nodes that are more active in G1 are shown in red, while those more active in G2 are shown in blue. Edges: The edges represent the interactions between nodes. The color of the edges indicates whether the interaction is stronger in G1 or G2. Interactions more prevalent in G1 are shown in magenta, and those stronger in G2 are shown in green.

This can provide valuable insights into the mechanisms driving the differential responses to the antibody treatment. the dysfunctional feedback loops in the non-responsive models may impede their ability to transduce the signal, resulting in a lack of clear treatment response.

These findings suggest that the structural integrity and composition of feedback loops are critical determinants of treatment efficacy. Focusing on these network characteristics can offer valuable insights on the molecular mechanisms underlying therapeutic responses and identify potential targets for enhancing treatment effectiveness in resistant cases.

### 1.3 Graph based Integrative approaches for patients’ classification and response prediction

To improve patient-specific treatment strategies, graph-based integrative approaches offer a powerful method for classifying patients and predicting their therapeutic responses. Using patient-specific gene regulatory networks, we employed a GCN to build a classifier capable of predicting responses to NRG1-Ab treatment.

Each PDX models personalized network was represented as a graph, with node features incorporating expression levels and attractor states at baseline. These comprehensive features enriched the graphs with key biological information. Using a population of networks per sample, we applied an 80-20% train-test split, resulting in 622 networks for training and 156 for testing. The GCN was trained to classify networks as either responsive or non-responsive based on their network structures and data-derived features, with treatment outcomes serving as labels.

The performance of the GCN model was evaluated using several metrics, including loss, accuracy, precision, recall, and balanced accuracy, along with cross-validation to ensure robustness. The loss over 3200 epochs showed a steady decrease for both the training and test sets, with the model achieving a train loss of 0.3736 and a test loss of 0.3630 at the final epoch. This consistent reduction in loss, along with the relatively small gap between training and test loss, indicates good generalization without significant overfitting (fig 4.A). Cross-validation yielded an average train recall of 0.9021 and test recall of 0.8610, along with an average train balanced accuracy of 0.9021 and test balanced accuracy of 0.8610, further confirming the models stability across different data splits.

**Fig 4.**
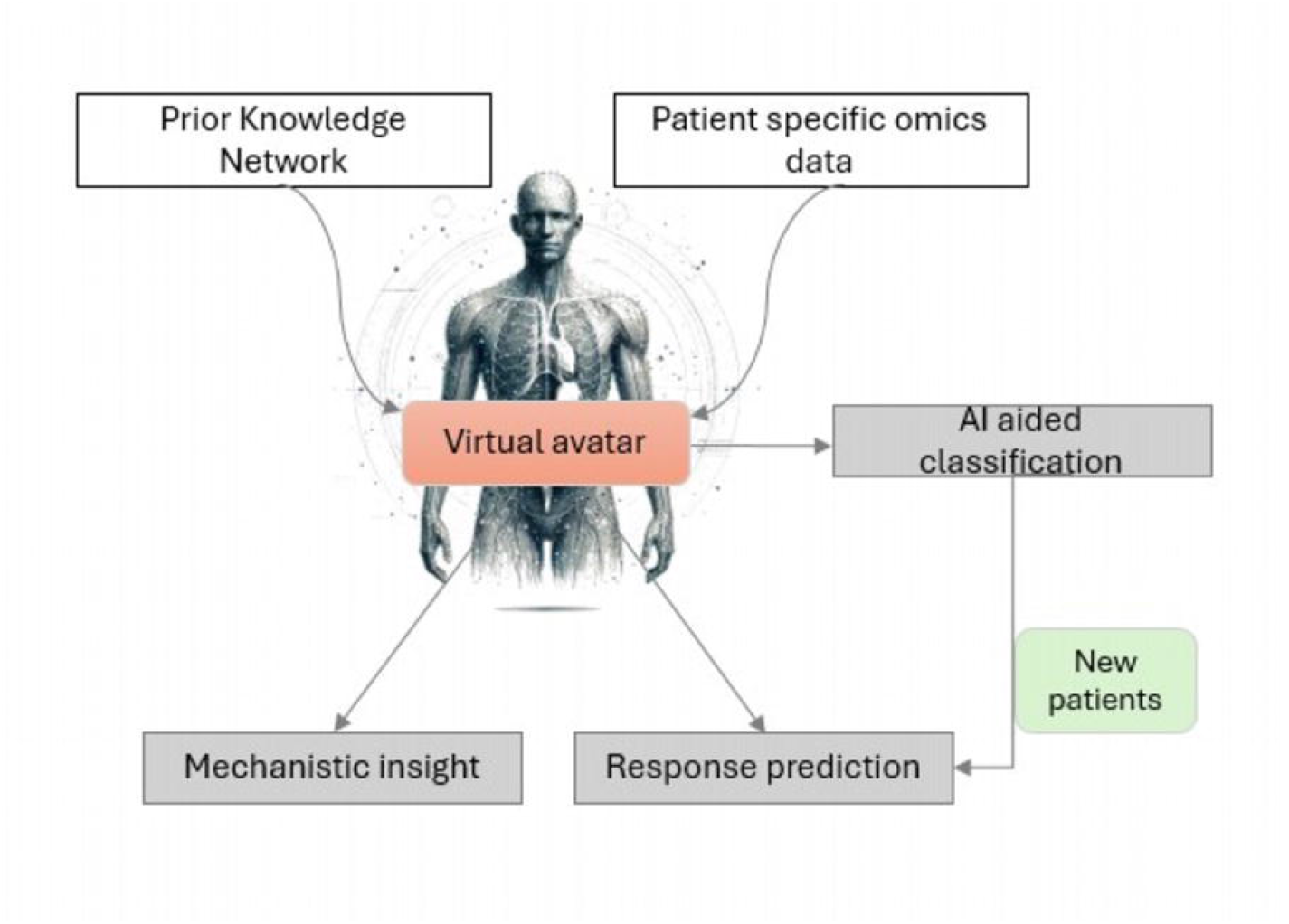
GCN model performance evaluation. (A) Loss Plot: The training and test loss curves over 3200 epochs. (B) ROC Curve: The Receiver Operating Characteristic (ROC) curve, showing the true positive rate versus the false positive rate for non-responsive and responsive classes. (C) Multidimensional Scaling (MDS) visualization of cosine distances between graph embeddings learned by the GCN. Points are colored by class (blue for non-responsive and orange for responsive)

The ROC curve further supports the model’s strong performance, with an AUC of 0.95 for both non-responsive (0) and responsive (1) classes, demonstrating the model’s high ability to distinguish between the two classes (fig 4.B). This suggests that the model has a strong discriminatory power across both classes, with a high true positive rate and a low false positive rate across various classification thresholds.

The Multidimensional Scaling (MDS) visualisation of cosine distances (fig 4.C) of the learned embeddings highlights the model’s ability to differentiate between responsive and non-responsive classes. While there is a clear separation between the two classes in the embedding space, some overlap remains, particularly between samples in the middle regions (fig 4.C). This overlap is likely responsible for the small drop in precision and recall, with the final test precision at 82.91% and test recall at 85.45 Overall, the GCN model demonstrates strong performance, achieving a test accuracy of 88.46% and balanced accuracy of 85.45%, with minimal signs of overfitting. The high AUC score, steady loss reduction, and reasonable separation in the MDS visualization of cosine distances of graph embeddings indicate that the model effectively learns the underlying structure of the data, but there remains room for improvement in distinguishing some ambiguous samples.

This GCN-based classifier provides a framework for predicting PDX responses to treatment using only baseline data. By leveraging personalized gene regulatory networks, this method allows for the integration of various biological data types into a single model, enhancing its predictive power. The ability to predict treatment outcomes based on baseline characteristics alone could facilitate more precise and personalized therapeutic interventions for future patients.

## 2 Proposed position and discussion

This study demonstrates the utility of graph-based integrative approaches for predicting patient responses to treatment, leveraging the power of personalized gene regulatory networks and GCNs. By integrating diverse biological data types, we can build models capable of predicting treatment efficacy with high accuracy, even from baseline data alone. These findings highlight the potential of patient derived models and GCNs as tools for advancing personalized medicine, offering a framework that could be applied across a wide range of therapeutic contexts.

In future, virtual avatars, created from patient-specific gene regulatory networks, will allow for highly personalized simulations of disease progression and treatment response. This will enable the design of therapeutic interventions tailored to the unique biological context of each patient.

AI plays a crucial role in enhancing the predictive power of these personalized models. By leveraging AI, we can integrate diverse types of biological data, such as gene expression, mutational profiles, and omics data, into graph-based models.

The potential for broader application of these technologies is immense, promising a new era of precision medicine where treatments are tailored to the unique genetic and molecular profiles of individual patients. By accurately predicting which patients will respond to specific treatments, healthcare providers can make more informed decisions, leading to better patient outcomes and reduced healthcare costs.

In addition to clinical applications, these technologies can accelerate the drug development process. By identifying likely responders early in clinical trials, drug developers can streamline the process of evaluating new therapeutics. This enhances the efficiency and success rate of new treatments, ensuring that only the most promising drugs advance through the costly and time-consuming stages of clinical development.

The methodologies demonstrated in our study are highly adaptable and can be applied across various diseases and treatment modalities. This scalability ensures that a wide range of patients can benefit from personalized therapeutic strategies.

## 3 Material and methods

### 3.1 Building executable patient derived models

#### 3.1.1 PKN Construction

We initiated the construction of the PKN from the hallmarks of cancer as outlined by Hanahan and Weinberg [20]. To identify the key genes associated with each hallmark, we retrieved the relevant gene sets from MSigDB [21]. Next, we used Omnipath [23], a comprehensive resource of curated signaling pathways, to interconnect these genes, forming a directed network. In this network, the edges between nodes represent the regulatory relationships, which are either inhibitory or activatory, based on the underlying biological interactions. Additionally, we incorporated information on complex protein components using the SIGNOR database [22]. In cases where interactions involved multi-component complexes, AND gates were employed to represent the need for simultaneous activation of multiple components for signal transduction. These logical gates allowed us to capture the cooperative nature of specific molecular events.

#### 3.1.2 Data binarization for integration into the logical model

To integrate continuous gene expression data into the logical model, the data must first be binarized into discrete states (0 or 1), representing inactive or active expression levels, respectively. For this study, RNA-seq data were binarized following the method described by Beal et al. [24]. Gene expression data are first classified in three categories according to their distribution across samples: bimodal, unimodal, and zero-inflated distribution. different distributions are then binarized differently as described by the authors.

#### 3.1.3 Model personalization using genetic algorithm

For each sample, we personalized the PKN by contextualizing it with the binarized gene expression data specific to each sample. This process was conducted using OptimusQual [18], an optimization tool designed to tailor the PKN to individual expression profiles.

The genetic algorithm employed iteratively adjusted the network to optimize its alignment with the profiles of the respective PDX sample. The optimization sought to maximize the score by prioritizing three key objectives: (1) maximizing the inclusion of essential nodes related to key cancer processes, such as Apoptosis, Proliferation, Metastasis, Hypoxia, Invasion-promoting macrophages, TGF*β* signaling, DNA repair, Angiogenesis, Immortality, Inflammation, Epithelial-mesenchymal transition (EMT), EGFR signaling, and NRG1 signaling; (2) minimizing the total number of nodes, the minimum being those present in the training set to maintain network simplicity; and (3) maximizing the number of edges to closely represent molecular interactions present in the PKN.

#### 3.1.4 Attractor state calculation

For each optimized sample-specific network, we calculated the attractor states using the mpbn (Most Permissive Boolean Networks) framework [25].

### 3.2 Personalized networks groups integration

Within-sample edge frequency was calculated by computing the mean edge frequency across all replicate networks optimized for each sample. Then, for each model and treatment combination, Within-group edge frequency *Fg* was calculated by computing the mean edge frequency across all samples belonging to that group.

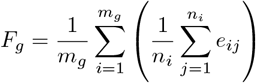

- *e*_*ij*_ represent the presence of edge *e* in replicate network *j* of sample *e* for a particular group.
- *n*_*i*_ represent the total number of replicate networks for sample *i*.
- *m*_*g*_ represent the total number of samples in group *g*.

For each common edge e present in both group g1 and group g2, the difference in edge weights is calculated as:

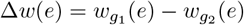

### 3.3 Graph classification

We used graph-structured data in this study, where each graph represented a personalized network. Nodes contained a 2-dimensional feature vector representing expression levels and attractor states (0 or 1) in a Boolean model. Each graph was classified into one of two target classes. The dataset was split into training and test sets using a stratified split, with a test size of 0.2 to maintain class balance.

A Graph Convolutional Network (GCN) was implemented using PyTorch and PyTorch Geometric. The model consisted of two graph convolutional layers, each with 64 output channels, followed by a ReLU activation and dropout (0.5) to prevent overfitting. A global mean pooling layer aggregated node embeddings into a single graph-level representation, which was passed to a fully connected layer for classification. Log-softmax was applied to compute class probabilities.

Class imbalance was addressed by computing class weights based on the inverse frequency of each class. The model was trained using the Adam optimizer with a learning rate of 0.001 and a weighted cross-entropy loss function. Training was performed for 3200 epochs with a batch size of 64.

Evaluation was conducted after each epoch on both the training and test sets, with the model in evaluation mode to disable dropout and batch normalization. Performance metrics included accuracy, precision, recall, and balanced accuracy.

To evaluate model robustness, we employed a three-fold stratified cross-validation on the combined training and test data, ensuring balanced representation of the target classes in each fold.

## Notes

### Competing Interest Statement

The authors have declared no competing interest.

## References

1. Eduati F, Jaaks P, Wappler J, Cramer T, Merten CA, Garnett MJ, et al. Patientspecific logic models of signaling pathways from screenings on cancer biopsies to prioritize personalized combination therapies. Molecular Systems Biology. 2020;16(2):e8664. doi:10.15252/msb.20188664.

2. Montagud A, Bal J, Tobalina L, Traynard P, Subramanian V, Szalai B, et al. Patient-specific Boolean models of signalling networks guide personalised treatments. eLife. 2022;11:e72626. doi:10.7554/eLife.72626.

3. Guex N, Crespo I, Bron S, Ifticene-Treboux A, Faes-vant Hull E, Kharoubi S, et al. Angiogenic Activity of Breast Cancer Patients Monocytes Reverted by Combined Use of Systems Modeling and Experimental Approaches. PLOS Computational Biology. 2015;11(3):e1004050. doi:10.1371/journal.pcbi.1004050.

4. Singh V, Naldi A, Soliman S, Niarakis A. A large-scale Boolean model of the rheumatoid arthritis fibroblast-like synoviocytes predicts drug synergies in the arthritic joint. npj Systems Biology and Applications. 2023;9(1):33. doi:10.1038/s41540-023-00294-5.

5. GarridoRodriguez M, Zirngibl K, Ivanova O, Lobentanzer S, SaezRodriguez J. Integrating knowledge and omics to decipher mechanisms via largescale models of signaling networks. Molecular Systems Biology. 2022;18(7):e11036. doi:10.15252/msb.202211036.

6. SaezRodriguez J, Blthgen N. Personalized signaling models for personalized treatments. Molecular Systems Biology. 2020;16(1):e9042. doi:10.15252/msb.20199042.

7. Papp O, Jordn V, Hetey S, Balzs R, Kaszs V, Bartha, et al. Network-driven cancer cell avatars for combination discovery and biomarker identification for DNA damage response inhibitors. npj Systems Biology and Applications. 2024;10(1):68. doi:10.1038/s41540-024-00394-w.

8. Kim D, Tran A, Kim HJ, Lin Y, Yang JYH, Yang P. Gene regulatory network reconstruction: harnessing the power of single-cell multi-omic data. npj Systems Biology and Applications. 2023;9(1):51. doi:10.1038/s41540-023-00312-6.

9. Muzio G, OBray L, Borgwardt K. Biological network analysis with deep learning. Briefings in Bioinformatics. 2021;22(2):1515–1530. doi:10.1093/bib/bbaa257.

10. Tsirvouli E, Nol V, Flobak Calzone L, Kuiper M. Dynamic Boolean modeling of molecular and cellular interactions in psoriasis predicts drug target candidates. iScience. 2024;27(2):108859. doi:10.1016/j.isci.2024.108859.

11. Singh N, Khan FM, Bala L, Vera J, Wolkenhauer O, Ptzer B, et al. Logic-based modeling and drug repurposing for the prediction of novel therapeutic targets and combination regimens against E2F1-driven melanoma progression. BMC Chemistry. 2023;17(1):161. doi:10.1186/s13065-023-01082-2.

12. Marbach D, Costello JC, Kffner R, Vega N, Prill RJ, Camacho DM, et al. Wisdom of crowds for robust gene network inference. Nature methods. 2012;9(8):796–804. doi:10.1038/nmeth.2016.

13. Wu S, Jin K, Tang M, Xia Y, Gao W. Inference of Gene Regulatory Networks Based on Multi-view Hierarchical Hypergraphs. Interdisciplinary Sciences: Computational Life Sciences. 2024;16(2):318–332. doi:10.1007/s12539-024-00604-3.

14. Wang T, Shao W, Huang Z, Tang H, Zhang J, Ding Z, et al. MOGONET integrates multi-omics data using graph convolutional networks allowing patient classification and biomarker identification. Nature Communications. 2021;12:3445. doi:10.1038/s41467-021-23774-w.

15. Wekesa JS, Kimwele M. A review of multi-omics data integration through deep learning approaches for disease diagnosis, prognosis, and treatment. Frontiers in Genetics. 2023;14. doi:10.3389/fgene.2023.1199087.

16. Roohani Y, Huang K, Leskovec J. Predicting transcriptional outcomes of novel multigene perturbations with GEARS. Nature Biotechnology. 2024;42(6):927–935. doi:10.1038/s41587-023-01905-6.

17. Li MM, Huang Y, Sumathipala M, Liang MQ, Valdeolivas A, Ananthakrishnan AN, et al. Contextual AI models for single-cell protein biology. Nature Methods. 2024;doi:10.1038/s41592-024-02341-3.

18. Dorier J, Crespo I, Niknejad A, Liechti R, Ebeling M, Xenarios I. Boolean regulatory network reconstruction using literature based knowledge with a genetic algorithm optimization method. BMC Bioinformatics. 2016;17(1):410. doi:10.1186/s12859-016-1287-z.

19. Kenichiro Ono KY, inventor; Medical; Biological Laboratories Co Ltd, assignees. Antibodies to human NRG1 protein. EP2955226A4; 2016. Available from: https://patents.google.com/patent/EP2955226A4/en.

20. Hanahan D. Hallmarks of Cancer: New Dimensions. Cancer Discovery. 2022;12(1):31–46. doi:10.1158/2159-8290.CD-21-1059.

21. Liberzon A, Birger C, Thorvaldsdttir H, Ghandi M, Mesirov J, Tamayo P. The Molecular Signatures Database Hallmark Gene Set Collection. Cell Systems. 2015;1(6):417–425. doi:10.1016/j.cels.2015.12.004.

22. LoSurdo P, Iannuccelli M, Contino S, Castagnoli L, Licata L, Cesareni G, et al. SIGNOR 3.0, the SIGnaling network open resource 3.0: 2022 update. Nucleic Acids Research. 2023;51(D1):D631–D637. doi:10.1093/nar/gkac883.

23. Trei D, Valdeolivas A, Gul L, PalacioEscat N, Klein M, Ivanova O, et al. Integrated intra and intercellular signaling knowledge for multicellular omics analysis. Molecular Systems Biology. 2021;17(3):e9923. doi:10.15252/msb.20209923.

24. Bal J, Montagud A, Traynard P, Barillot E, Calzone L. Personalization of Logical Models With Multi-Omics Data Allows Clinical Stratification of Patients. Frontiers in Physiology. 2019;9:1965. doi:10.3389/fphys.2018.01965.

25. Paulev L, Kolk J, Chatain T, Haar S. Reconciling qualitative, abstract, and scalable modeling of biological networks. Nature Communications. 2020;11(1):4256. doi:10.1038/s41467-020-18112-5.

